# Eukaryotic composition across seasons and social groups in the gut microbiota of wild baboons

**DOI:** 10.1101/2024.12.17.628920

**Authors:** Mary Chege, Pamela Ferretti, Shasta Webb, Rosaline W. Macharia, George Obiero, Joseph Kamau, Susan C. Alberts, Jenny Tung, Mercy Y. Akinyi, Elizabeth A Archie

**Affiliations:** One Health Centre, Kenya Institute of Primate Research, Nairobi, Kenya; Department of Biochemistry, University of Nairobi, Nairobi, Kenya; Department of Biological Sciences, University of Notre Dame, Notre Dame, Indiana, USA; Department of Medicine, Genetic Medicine Section, University of Chicago, Chicago, USA; Departments of Biology and Evolutionary Anthropology, Duke University, Durham, NC, USA; Duke University Population Research Institute, Duke University, Durham, NC, USA; Department of Primate Behavior and Evolution, Max Planck Institute for Evolutionary Anthropology, 04103 Leipzig, Germany; Canadian Institute for Advanced Research, Toronto, Ontario, Canada

**Author notes:** Corresponding authors: Mary Chege Elizabeth Archie.

**Keywords:** eukaryotes, gut microbiome, wild baboons, fungi, protists, social groups

## Abstract

**Background:** Animals coexist with complex microbiota, including bacteria, viruses, and eukaryotes (e.g., fungi, protists, and helminths). While the composition of bacterial and viral components of animal microbiota are increasingly well understood, eukaryotic composition remains neglected. Here we characterized eukaryotic diversity in the microbiomes in wild baboons and tested the degree to which eukaryotic community composition was predicted by host social group membership, sex, age, and season of sample collection.

**Results:** We analyzed a total of 75 fecal samples collected between 2012 and 2014 from 73 wild baboons in the Amboseli ecosystem in Kenya. DNA from these samples was subjected to shotgun metagenomic sequencing, revealing members of the kingdoms Protista, Chromista, and Fungi in 90.7%, 46.7%, and 20.3% of samples, respectively. Social group membership explained 11.2% of the global diversity in gut eukaryotic species composition, but we did not detect statistically significant effect of season, host age, and host sex. Across samples, the most prevalent protists were *Entamoeba coli* (74.66% of samples), *Enteromonas hominis* (53.33% of samples), and *Blastocystis subtype* 3 (38.66% of samples), while the most prevalent fungi included *Pichia manshurica* (14.66% of samples), and *Ogataea naganishii* (6.66% of samples).

**Conclusions:** Protista, Chromista, and Fungi are common members of the gut microbiome of wild baboons. More work on eukaryotic members of primate gut microbiota is essential for primate health monitoring and management strategies.

## Introduction

Vertebrate gut microbiota are complex and dynamic communities of bacteria, viruses, archaea, and eukaryotes [1,2]. To date, most research on vertebrate gut microbiota has focused on bacteria, in part because bacteria are both easy to characterize via 16S rRNA gene sequencing [3,4] and because bacteria are important to host health, with well-known effects on host metabolism, vitamin biosynthesis, and immune modulation [5–8]. However, vertebrate gut microbiota also contain eukaryotes, such as protists, metazoans, and fungi. Research on these eukaryotic communities remains neglected, in part because genetic and bioinformatic methods to characterize these communities are less developed than those for bacteria [9–12].

To date, the best characterized eukaryotic gut communities are those found in humans and laboratory mice. Humans living in a wide range of geographic locations harbor gut eukaryotes, including protists such as *Blastocystis*, *Entamoeba*, and *Enteromonas,* and fungi such as *Saccharomyces*, *Candida*, *Penicillium*, *Aspergillus*, and *Malassezia* [13–15]. The relevance of these taxa to host health often remains unclear [16]. Some of these protists, such as *Entamoeba histolytica*, *Blastocystis hominis, and Aspergillus fumigatus,* may have pathogenic effects on hosts [14,17–21], while many others likely have commensal relationships with their hosts, and their presence may indicate a normal, healthy gut microbiota [16,22,23]. For instance, in mice, *Tritrichomonas musculis* is associated with host immune modulation and protection against bacterial mucosal infections [24].

In contrast, little is known about the eukaryotes living in the intestines of wild animals, including non-human primates [25–27]. Here we characterize eukaryotic members of the gut microbiota of wild baboons (*Papio sp.*) living in the Amboseli ecosystem in Kenya [28]. Prior work in this population has revealed several variables that influence individual exposure and susceptibility to gut bacteria, and we predicted that these same variables would also be important in predicting gut eukaryotic composition, including host social group membership [29–32], the season of sample collection [33–39], and host age [34,40–42]. For instance, social group membership influences baboon ranging patterns, resource use, and social relationships, which might influence microorganism exposure and transmission. In support, baboons from different social groups show distinct gut bacterial communities, and social group membership explains more variance in the gut bacterial microbiome than host age or sex [29]. Seasonality also shapes baboon diet, water source use, and other aspects of the environment, leading to systematic fluctuations in the abundance of several bacterial taxa as a function of the season of sample collection [43].

Our objectives in this paper were to characterize common gut eukaryotes in wild baboons and test the effects of seasonality, host social group membership, host sex, and host age on baboon gut eukaryotic community composition and diversity. We accomplished this objective by leveraging two shotgun metagenomic data sets: (i) samples from 48 individual baboons living in two social groups in July and August 2012 [29], and (ii) samples from 27 individual baboons collected during the wet (April and May) and dry seasons (September) of 2014 (two individuals were included in both data sets so the total sample size was 75 samples from 73 individual baboons). We expected that gut eukaryotic communities would be influenced by similar factors to gut bacterial communities in this population. Our results provide new insights into the gut eukaryotic composition of wild baboons, contributing useful information for understanding the biology and health of wild primates.

## Materials and methods

### Study subjects and sample material

The baboons sampled in this study were studied by the Amboseli Baboon Research Project (ABRP) in the Amboseli ecosystem in Kenya. Founded in 1971, the ABRP conducts longitudinal research on known individual baboons living in several social groups [28]. Members of this population are hybrids between yellow and anubis baboons (*Papio cynocephalus* and *P. anubis*) [44–46]. The Amboseli ecosystem experiences a 5-month dry season from June through October, followed by a 7-month wet season with highly variable rainfall [47].

Data on the Amboseli baboons is collected by experienced observers year-round during 5-hour monitoring sessions, six days per week. During these sessions, observers collect fecal samples opportunistically from individuals, all of which are known to the observers from distinctive physical features. For each sample, the baboon’s social group membership is known from group censuses collected during each monitoring session. Age is known to within a few days’ error for all individuals born into ABRP study groups (n=64 baboons in this study), and estimated for other individuals based on observable morphological characteristics and body condition (n=9 baboons who were all immigrant males, as males but not females disperse from their natal groups in this population). Sex is known based on morphological differences. All sampled individuals appeared healthy upon visual examination. All fecal samples are collected within a few minutes of defecation, thoroughly mixed, and preserved in 95% ethanol (2:5 feces to ethanol).

We analyzed eukaryotic composition in two sets of fecal samples. The first set was used to test for social group effects on eukaryotic composition. This set included samples from 48 adult baboons living in two social groups: ‘Mica’s group’, (11 females and 8 males), and ‘Viola’s group’, (20 females and 9 males). These samples were collected within a one-month window during the dry season of 2012 (**Figure 1**; **Supplementary Table 1;** an analysis of bacterial microbiome composition from these samples was published in Tung et al. [29]). The second dataset was used to test for seasonal differences in eukaryotic composition. It included samples from 27 baboons collected during the dry season in September 2014 (n=15) and the wet season between April and May 2014 (n=12) (**Figure 1; Supplementary Table 1**). Host sex and age were known for individuals in both data sets and two individuals occurred in both data sets.

**Figure 1.**
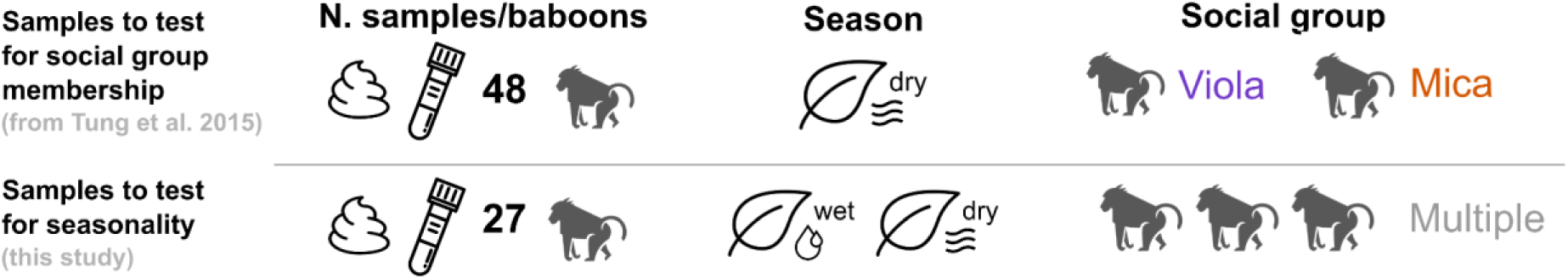
Schematic of the two sets of fecal samples investigated in this study (**Supplementary Table 1**). Shotgun metagenomic data for the first set of samples on social group membership was published in 2015 by Tung et al. [29]; these data are publicly available in the NCBI’s Short Read Archive (Bioproject PRJNA271618). Shotgun metagenomic data for the second set of samples on seasonality were generated for the present study; these data are publicly available on NCBI’s Short Read Archive (Bioproject PRJEB81717). Data on host sex and age were available for samples in both data sets.

### Genomic DNA extraction and sequencing

Total DNA was extracted from the first (social group) set of fecal samples using the MO BIO Laboratories, Inc., Carlsbad, CA, PowerSoil DNA Isolation kit. Samples for the second (seasonal) set were extracted using Qiagen’s DNeasy PowerSoil kit (Venlo, Netherlands). Both protocols were performed according to the manufacturer’s instructions.

For samples in the first (social group) dataset [29], 200 ng of DNA were prepared for metagenomic sequencing on an Illumina HiSeq 2500, using Kapa Biosystems Library Preparation Kits (Kapa Biosystems, Wilmington, MA). The DNA samples were sheared to an average size of 400 base pairs, followed by ligation to barcoded adapters. The libraries were subjected to 100 base pair paired-end sequencing at the UCLA Neuroscience Genomics Core. In total, 1.4 billion raw, paired-end Illumina sequences were generated, with a mean ± SD of 14.4 ± 13.7 million read pairs per sample. The raw metagenomic sequencing data are deposited in the NCBI’s Short Read Archive (BioProject PRJNA271618).

For samples in the second (seasonal) dataset, 200 ng of DNA were prepared for metagenomic sequencing on the Illumina NovaSeq X, utilizing the SeqWell purePlex DNA Library Preparation Kit (SeqWell, Beverly, MA). Samples were subjected to 150 base pair paired-end sequencing at the University of Minnesota Genomics Core. In total, 2.54 billion raw, paired-end Illumina sequences were generated, with a mean ± SD of 94 ± 31.2 million read pairs per sample. All the raw reads for the second dataset (n = 27) are deposited in NCBI’s Short Read Archive (BioProject PRJEB81717).

### Identifying dominant eukaryotic members of the baboon gut microbiome

The raw data were processed using FastQC [48] to assess the quality of the reads. Duplicate reads were removed using FastUniq [49] and trimmed to remove adapter sequences and low-quality bases (PHRED score <20) using Trimmomatic (v0.39) in paired-end mode [50]. We also removed read pairs where one read was shorter than 75 base pairs after trimming.

We then used two previously-developed pipelines to further filter our sequences and identify the presence/absence of eukaryotic taxa in each sample: EukDetect [51] (https://github.com/allind/EukDetect) and its descendent pipeline gutprotist-search https://github.com/allind/gutprotist-search). In brief, EukDetect aligns reads to a database consisting of conserved eukaryotic marker genes from curated whole genome assemblies [51]. Gutprotist-search, developed alongside EukDetect, complements the approach in EukDetect by using a database of NCBI sequences for particular taxonomic identifiers for eukaryotic taxa that lack genome assemblies. Because genome sequences are unavailable for many gut eukaryotes, gutprotist-search helps identify taxa that might otherwise go undetected.

Both pipelines were run using the recommended Snakemake workflow engine [51,52] with default parameters. The metagenomic reads were aligned to the EukDetect marker database and the gutprotist-search database using Bowtie2 [53], followed by stringent quality filtering based on mapping quality to retain reads with a PHRED score ≥30, ensuring high base-call accuracy. Sequence complexity was assessed using a complexity score threshold of ≥0.5 to retain only high-complexity reads, reducing spurious alignments due to low-complexity regions. Additional filtering as described in EukDetect protocol extended these steps to refine taxonomic assignments by addressing off-target alignments arising from false positives. Specifically, reads mapping to multiple species within a genus were compared for sequence identity and coverage, with lower confidence species excluded, and the ETE3 toolkit [54] was used to validate alignment accuracy. Following recommended practice, only taxa with more than four reads aligning to at least two distinct marker genes were retained, but unfiltered results were also explored for low-abundance species as recommended by the authors [51]. The EukDetect and gutprotist-search pipelines output a list of the eukaryotic taxa identified in the samples and associated relevant statistics such as the number of observed marker genes per sample per taxon, number of reads mapping to the marker genes, total marker gene coverage and identity percentage. We combined the results of these pipelines for each sample into a single table for our analyses (**Supplementary Table 2**). In this table, taxa were marked present (1) in a given sample if they were detected in one or both pipelines and absent (0) from a sample if they were not detected by either pipeline. Finally, the insect metazoan *Callosobruchus maculatus* was detected in 1 sample, but excluded from downstream analyses as it was likely acquired from food, and therefore not part of the gut eukaryotic community.

### Statistical analyses

All statistical analyses were conducted in the R statistical environment (R version 4.3.3) [55]. We began our analyses by reporting the eukaryotic taxa that we identified across both sets of samples (N=75 samples; 48 from the first (social) set and 27 from the second (seasonal) set).

Next, to test whether eukaryotic species richness differed between baboon social groups (first sample set) or season (second sample set), we calculated the Simpson’s alpha diversity index using the package *vegan* [56]. We then tested for differences in alpha diversity as a function of social group, season, host age, host sex, and the number of paired-end sequences generated for each sample using a linear model implemented in the *stats* package [55].

To test for compositional differences between eukaryotes as a function of baboon social group or season, we calculated Jaccard distance matrices within each data set using the vegdist function in *vegan* [56] (Jaccard distances measure dissimilarity between samples based on the presence/absence of taxa in each sample). To visualize how eukaryotic species composition varied between social groups or seasons, we used non-metric multidimensional scaling (NMDS). To determine the proportion of variance in community composition attributable to social group or season and host sex or age, we conducted a permutational multivariate analysis of variance (PERMANOVA) implemented in the adonis function in *vegan*, with 10,000 permutations [56] To test the effect of social group and season on the presence of each eukaryotic species within each data set using a binomial generalized linear model (GLM) implemented in the *stats* package. The presence or absence of each eukaryotic species was modeled as a response variable while the predictor variables were social group membership (Mica’s or Viola’s), season (wet or dry), host sex, host age, and read count.

### Ethics statement

All protocols were approved by the Institutional Animal Care and Use Committee (IACUC) at the University of Notre Dame to cover behavior observations and fecal sample collections in baboons at Amboseli, under protocol number 22-05-7259.

## Results

### Metagenomic analysis reveals diverse gut eukaryotic communities in wild baboons

Across both the social (n=48) and seasonal (n=27) data sets, we found genetic evidence for 21 eukaryotic species (**Supplementary** Figure 1**; Supplementary Table 3;** range=0 to 7 taxa per sample). The mean number of eukaryotic taxa per sample across both data sets was 3.10 (SD= 1.60). Within each dataset, the average number of eukaryotic taxa present per sample was 3.09 (SD=1.47) in the social group dataset and 3.12 (SD= 1.84) in the seasonal dataset. Information on the potential transmission mode and health relevance of each taxon is in **Supplementary Table 4**. Additionally, we detected genetic evidence for the metazoan arthropod, *Callosobruchus maculatus* in 1.33% (n=1 of samples). Because baboons frequently ingest insects, these sequences may come from a closely-related insect in the baboon diet, and are not living members of the microbiome community. Sequences attributed to *C. maculatus* are removed from our analyses.

Overall, Protista was the most well-represented kingdom (n=68 samples; 90.7%), followed by Chromista (n=35 samples; 46.7%), and Fungi (n=22 samples; 29.3%; **Supplementary** Figure 1). Over half of the samples in the data set (n=56 samples; 74.7%) contained at least two detectable eukaryotic taxa, while 2.7% (n=2 samples) contained 7 eukaryotic taxa, the maximum number we observed. Only 2 samples (2.67%) had no detectable eukaryotes (one in the social group data set and one in the seasonal data set). Across all samples, the five most prevalent species were *Entamoeba coli* (n=56 samples; 74.7%)*, Enteromonas hominis* (n=40 samples; 53.3%)*, Blastocystis subtype 3* (n=29 samples; 38.7%)*, Iodamoeba sp* (n=27 samples; 36%), and *Chilomastix mensnili* (n=26 samples; 34.7%; **Supplementary Table 3**). We also detected a set of species found in one or only a handful of samples, including (*Candida blattae, Malassezia restricta*, *Ogatea naganashii, Preussia sp.,* and *Aspergillus sydowii* (see **Supplementary Table 3**).

### Social group membership predicts gut eukaryotic diversity and composition

In the first (social group membership) data set, we found significant differences in the gut eukaryotic alpha diversity for baboons in different social groups (linear model test, p=0.015;

**Figure 2A; Supplementary Table 5**). Samples from baboons living in Viola’s group exhibited higher Simpson’s diversity compared to those living in Mica’s group (**Figure 2A**).

**Figure 2.**
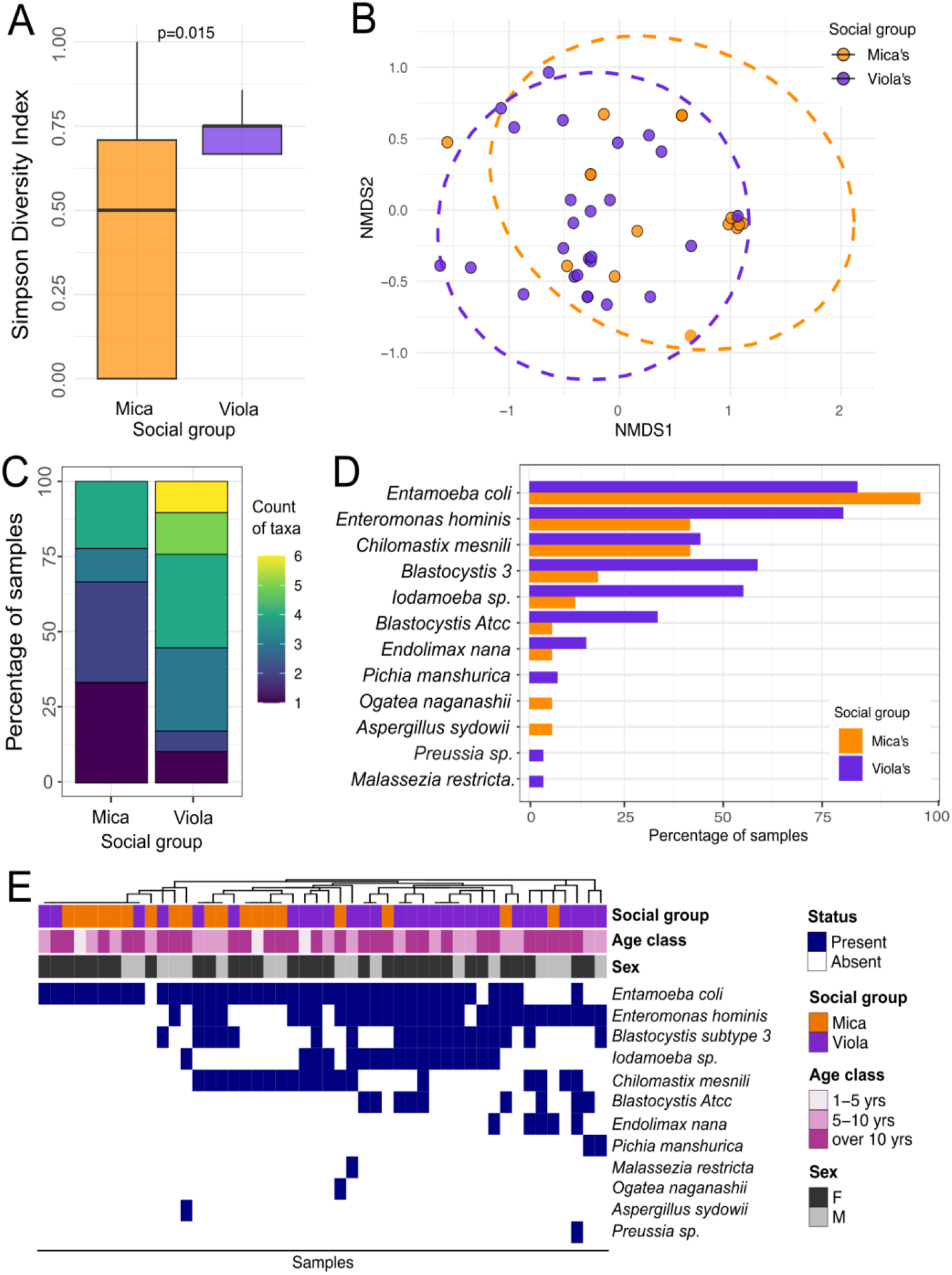
Social group membership is correlated with baboon gut eukaryotic composition. (A) Eukaryotic species diversity between Viola’s and Mica’s social groups using the Simpson’s diversity index. A linear model was used to calculate statistical significance. (B) Non-metric multidimensional scaling (NMDS) plot of gut eukaryotic composition as measured using Jaccard index dissimilarity matrices. (C) Number of eukaryotic taxa per sample in Mica and Viola’s social groups and (D) prevalence of eukaryotic taxa in the two social groups. (E) Heatmap of the eukaryotic taxonomic composition of samples used to test for social group membership, with each column representing one fecal sample and metadata on social group, season, age class, and sex of the host. Samples are clustered using Euclidean distance of eukaryotic community composition.

Eukaryotic composition was also significantly different between Viola’s and Mica’s groups, explaining 11.2% of the variation in eukaryotic composition (PERMANOVA: r^2^ = 0.111 p = 1×10^-3^; **Figure 2B; Supplementary Table 6**). We also found a trend such that the number of paired end sequences in each sample had a small effect on eukaryotic composition (r^2^ = 0.045; p = 0.05). By contrast, host age and sex did not make statistically significant contributions to gut eukaryotic composition (age, p = 0.45; sex, p = 0.504; see **Supplementary Table 6)**.

On average, 50% of Viola’s and 12.5% of Mica’s samples had at least 3 eukaryotic taxa (**Figure 2C**). In support of the patterns of alpha diversity we observed (**Figure 3A**), most taxa were more prevalent in Viola’s group as opposed to Mica’s group, including *Blastocystis subtype* 3 (coefficient: −2.03, p=0.01; **Supplementary Table 7**), *Blastocystis. ATCC* (coefficient: −1.99, p=0.08), *Enteromonas hominis* (coefficient: −1.77, p=0.01)*, Iodamoeba sp.* (coefficient: −2.21, p=0.01) and *Endolimax nana* (coefficient: −1.82, p=0.20). Only one taxon, *Entamoeba coli,* was more prevalent in the Mica’s group compared to the Viola’s group (**Figure 2D-E**). *Ogataea naganishii* and *Aspergillus sydowii* were found at low prevalence (2%) only in samples from Mica’s group, while *Preussia sp. BSL10* and *Malassezia restricta* were exclusively found in samples from Viola’s group (2%, **Figure 2D-E**).

**Figure 3.**
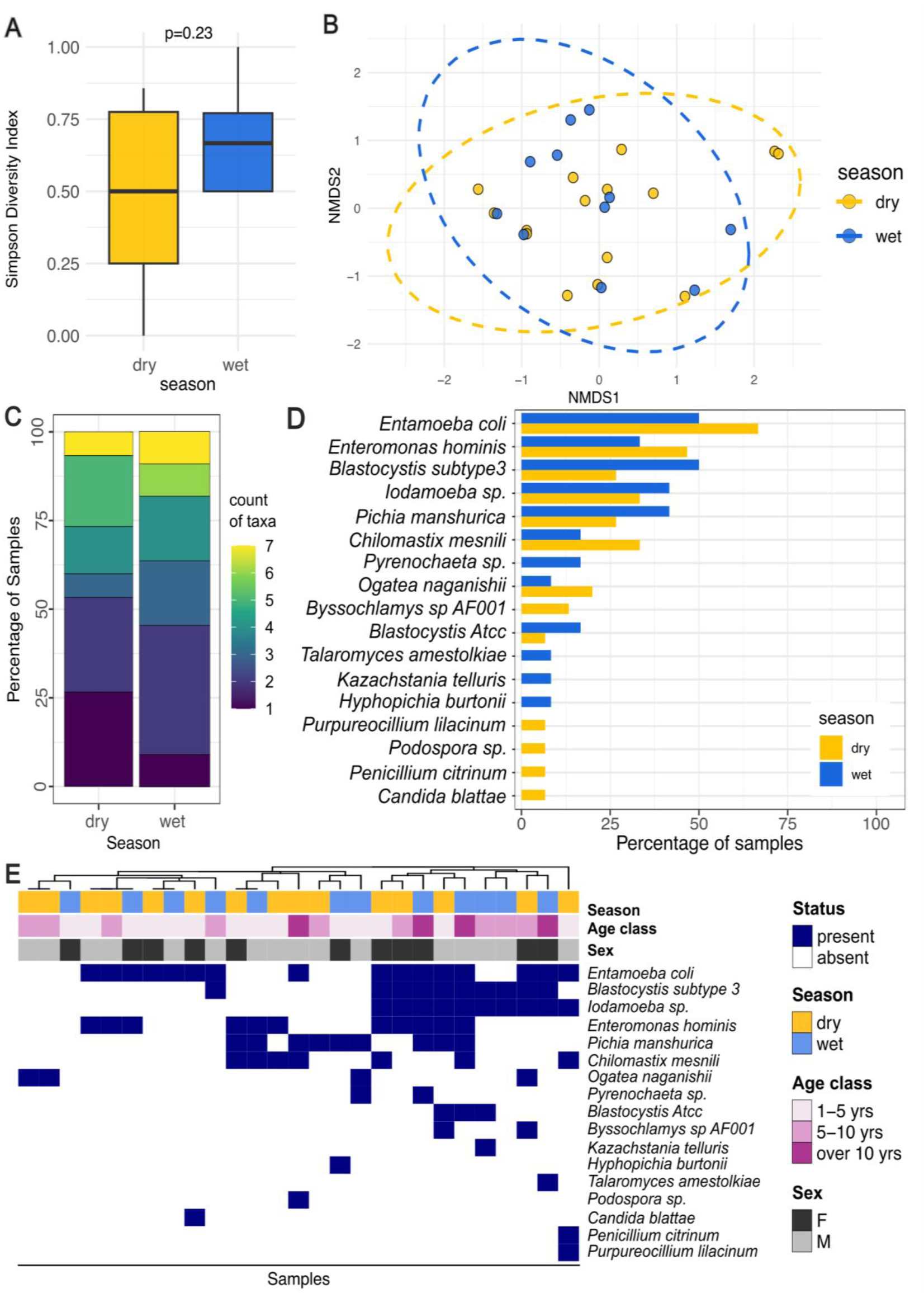
Season did not predict baboon gut eukaryotic diversity and composition. (A) Simpson’s diversity index of baboon gut microbiome composition between the wet and dry seasons. A linear model was used to calculate statistical significance. (B) Non-metric multidimensional scaling (NMDS) plot of gut eukaryotic microbiome composition as measured using Jaccard index dissimilarity matrices. (C) Number of eukaryotic taxa per sample in the dry and wet season. (D) prevalence of eukaryotic taxa in the wet and dry seasons. (E) Heatmap of the eukaryotic taxonomic composition of samples used to test for seasonality, with metadata on social group, season, age class, and sex of the host. Each column represents one fecal sample; samples are clustered using Euclidean distance of eukaryotic community composition.

### Season did not predict gut eukaryotic diversity and composition

We found no significant differences in the alpha diversity of the eukaryotes between samples collected in the dry season compared to the wet season (Simpson’s diversity: effect size = 3.6%, Linear model test p = 0.23; **Figure 3A**). Across all 27 samples in the seasonal dataset, no significant variation in taxonomic composition was explained by season, age, sex, or number of paired end sequences (PERMANOVA, r^2^ = 0.036; p = 0.48; age, p = 0.20; sex, p = 0. 64, number of paired end sequences, p = 0.13; **Figure 3B**).

On average, 25.9% of samples collected in the dry season and 22.2% of samples collected in the wet season contained at least 3 detectable eukaryotic taxa (**Figure 3C**). The eukaryotic communities across all the samples in the seasonal data set were clustered according to their diversity and composition, which indicated small taxon-level distinctions between the wet and the dry seasons, despite no strong evidence for global seasonal shifts. No species differed in prevalence between seasons (**Supplementary Table 7**). In line with our results on the social group dataset, *Entamoeba coli* emerged as the most prevalent and dominant species (n= 16 samples; 59.3%), followed by *Enteromonas hominis* (n= 11 samples; 40.7%), *Blastocystis* subtype 3 and *Iodamoeba* sp. (n= 10 samples each; 37% each), *Pichia manshurica* (n= 9 samples; 33.3%), *Chilomastix mesnili* (n= 7 samples; 25.9%), *Ogatea naganishii* (n= 4 samples; 14.8%), *Blastocystis* ATCC (n= 3 samples; 11.1%) and *Byssochlamys* sp. AF001 (n= 2 samples; 7.4%; **Figure 3D-E**).

## Discussion

Although eukaryotes are commonly found in mammalian gut microbiomes, little is known about their importance for host health or the factors that drive their prevalence in wild non-human primates. Here, we investigated the composition and diversity of the eukaryotic communities inhabiting the gastrointestinal tract of wild baboons, focusing on the explanatory power of host social group membership, season, host sex, and host age. We detected genetic evidence for several eukaryotic taxa, spanning 3 kingdoms (Fungi, Protista, and Chromista in order of prevalence) and 21 species. Most of the taxa we found are considered non-pathogenic commensals and have been previously found in human populations across the world [57]. Among primates, *Entamoeba coli* has been identified in mountain gorillas (*Gorilla beringei beringei*) [58], western lowland gorillas (*Gorilla gorilla gorilla*) [59], red colobus (*Procolobus badius tephrosceles*) [60,61], red-tailed monkeys (*Cercopithecus ascanius schmidti*) [60,61], vervet monkeys (*Cercopithecus aethiops pygerythrus*) [60,61], baboons (*Papio anubis*), and chimpanzees [60,61]. The high prevalence of *Entamoeba coli* among non-human primates might be attributed to their social behavior and communal living, as Entamoeba species are known to be transmitted by intake of a mature cyst through either ingestion of contaminated water or food, or direct oral-fecal contact [62,63], in addition to the commensal relationship between the parasite and its hosts.

Among the most abundant taxa identified in baboons was *Blastocystis* (56%). *Blastocystis* is a common chromista found in humans, and non-human primates such as baboons. gorillas, chimpanzees, and other mammals, as well as in birds [64–66]. Whether *Blastocystis* is commensal or pathogenic is still under debate [17]. In humans, members of the *Blastocystis* genus are common and stable colonizers of the gut of healthy individuals [67,68]. However, *Blastocystis* has also been associated with some gastrointestinal disorders such as inflammatory bowel disease (IBD) and irritable bowel syndrome (IBS) in humans [69], indicating possible pathogenicity. The role of *Blastocystis* in non-human primates remains unclear [66]. The high prevalence of *Blastocystis* among the baboons included in this study, which all appeared healthy upon visual examination when sampled, suggests that *Blastocystis* might be a commensal member of the gut microbiome of wild baboons.

The most common *Blastocystis* identified in the gut of the baboons in our study was *Blastocystis* subtype 3, found in 38.7% of the samples. *Blastocystis* subtype 3 has been previously found in humans and livestock [64], and is among the *Blastocystis* subtypes previously connected to human gut colonization [64,70]. Together, these results indicate the possibility for transmission of *Blastocystis* between wild baboons, livestock, and human populations, though further research is warranted to test this possibility. Transmission between species would not be surprising as both wild baboons and livestock share water holes during the dry season, and *Blastocystis* are commonly transmitted through shared water sources. Other taxa were less common in our samples (found in less than 10.7% of samples), and included *Candida blattae, Malassezia restricta* (which is a member of the normal primate skin microbiota, occasionally contributing to skin conditions), *Ogatea naganashii, Preussia sp.* (a non-pathogenic genus), and *Aspergillus sydowii* (which can cause respiratory infections in immunocompromised individuals; **Supplementary Table 4**). The latter was exclusively found in Mica’s group at low prevalence (2%), suggesting that, in contrast to recent observations in wild macaques (*Tibetan macaque*, *Macaca fascicularis,* and *Macaca namestrina*) [71,72], *Aspergillus* is not a common member of the wild baboon gut microbiome.

Among the variables we investigated, social group membership was the strongest predictor of eukaryotic diversity, explaining approximately 14.8% of the variation in eukaryotic community composition. Specifically, the larger social group (Viola) was characterized by greater eukaryotic diversity. This pattern may be due to the fact that individuals in larger groups interact with more hosts in larger as compared to smaller groups, and as a result they may be exposed to more diverse eukaryote communities. This result is consistent with a previous study from the Amboseli ecosystem [73] and another on geladas [74], showing that members of a larger social group exhibited higher diversity in their gut microbiota than individuals belonging to smaller groups. However, another possibility is that the increased eukaryotic diversity observed in Viola’s group is related to their home range occupancy. Viola’s home range was bigger than Mica’s at the time the samples we analyzed were collected [29], suggesting that Viola’s group could have been exposed to a wider variety of resources, substrates, and, ultimately, microbes. Other variables such as sex and age were less relevant in explaining eukaryotic diversity in our sample (1.6 % and 1.7%, respectively). Combined, these results suggest that social group membership plays an important role in eukaryotes’ ability to colonize hosts, and are in line with previous studies that reported a clear association between social group membership and the composition of bacterial communities in the gut of humans, non-human primates, carnivores, rodents, insects, and birds [29,75–83]. Physical contact, shared environment, and group behaviors might therefore influence both prokaryotic and eukaryotic diversity and composition via similar mechanisms.

We found that eukaryotic diversity was slightly higher during the wet season compared to the dry season, and that the eukaryotic composition in the gut varied across seasons. However, these trends were not large nor statistically significant. This result contrasts with previous studies that identified seasonality as highly relevant in shaping the bacterial component of the gut microbiome of human [37] and animal populations [33,35,36], including baboons [35–37,84]. However, this result may be due to the limited sample size of the second dataset used to investigate the impact of seasonality. Further investigation in larger cohorts is warranted to assess whether seasonality significantly impacts eukaryotic diversity in the gut microbiome of wild baboons. An additional limitation of our study is that the analysis of season on gut microbiome eukaryotic composition was performed on a cohort with highly heterogeneous group membership, and we could not use them to analyze the effects of social group membership due to low statistical power for this variable. Likewise, the social group membership sample set was collected entirely in the dry season and could not be used to assess seasonal variability. Therefore, the importance of these two variables in relation to each other is not clear from this study.

### Conclusions

To our knowledge, this is the first metagenomic study to characterize the eukaryotic gut microbiota of wild baboons. Taken together, our results indicate that eukaryotes are an important part of the microbial communities inhabiting the gut of wild baboons, and that social group membership plays a role in shaping gut eukaryotic composition and diversity over time. Understanding how social factors affect microbiome composition could therefore be informative about the evolution of social behavior and its health implications, and reinforces the importance of considering social dynamics in microbiome research.

## Data availability

The raw metagenomic sequences for the social data set dataset and the seasonal data set presented in this study are available on NCBI Sequence Read Archive (SRA) under the BioProject accession numbers PRJNA271618 and PRJEB81717, respectively. Comprehensive metadata for the samples introduced in this study are available as the Supplementary Material. Code is available on github: https://github.com/ArchieLab/chege_etal_2024.

## Acknowledgements

We thank Jeanne Altmann for her essential role in stewarding the Amboseli Baboon Project. In Kenya, we thank the Kenya Wildlife Service, the Wildlife Training and Research Institute, the National Council for Science, Technology, and Innovation, and the National Environment Management Authority for permission to conduct research and collect biological samples. We also thank the University of Nairobi, Kenya Institute of Primate Research, National Museums of Kenya, the Amboseli-Longido pastoralist communities, the Enduimet Wildlife Management Area, Ker & Downey Safaris, Air Kenya, and Safarilink for their cooperation and assistance in the field. We thank Karl Pinc for managing and designing the database. We thank Raphael Mututua, Serah Sayialel, Kinyua Warutere, Lilian Musembei, and Long’ida Siodi for collecting the field data and fecal samples. Complete acknowledgments for the Amboseli Baboon Research Project can be found online at https://amboselibaboons.nd.edu/acknowledgements/.

## Author contributions

Conceptualization, EAA, MC, MYA, JT; Sample and metadata collection, EAA, JT, SCA; Samples processing and data analysis, MC, SW, PF; Writing – original draft, MC.; Writing – review & editing, MC, PF, SW, EAA, MYA, RM, GO, JK, JT, SCA; Supervision, RWM, GO, MYA, EAA; Funding acquisition, EAA, JT, SCA. All authors read and approved the final manuscript.

