## Supplementary Figures for "Eukaryotic composition across seasons and social groups in the gut microbiota of wild baboons"

†corresponding authors:

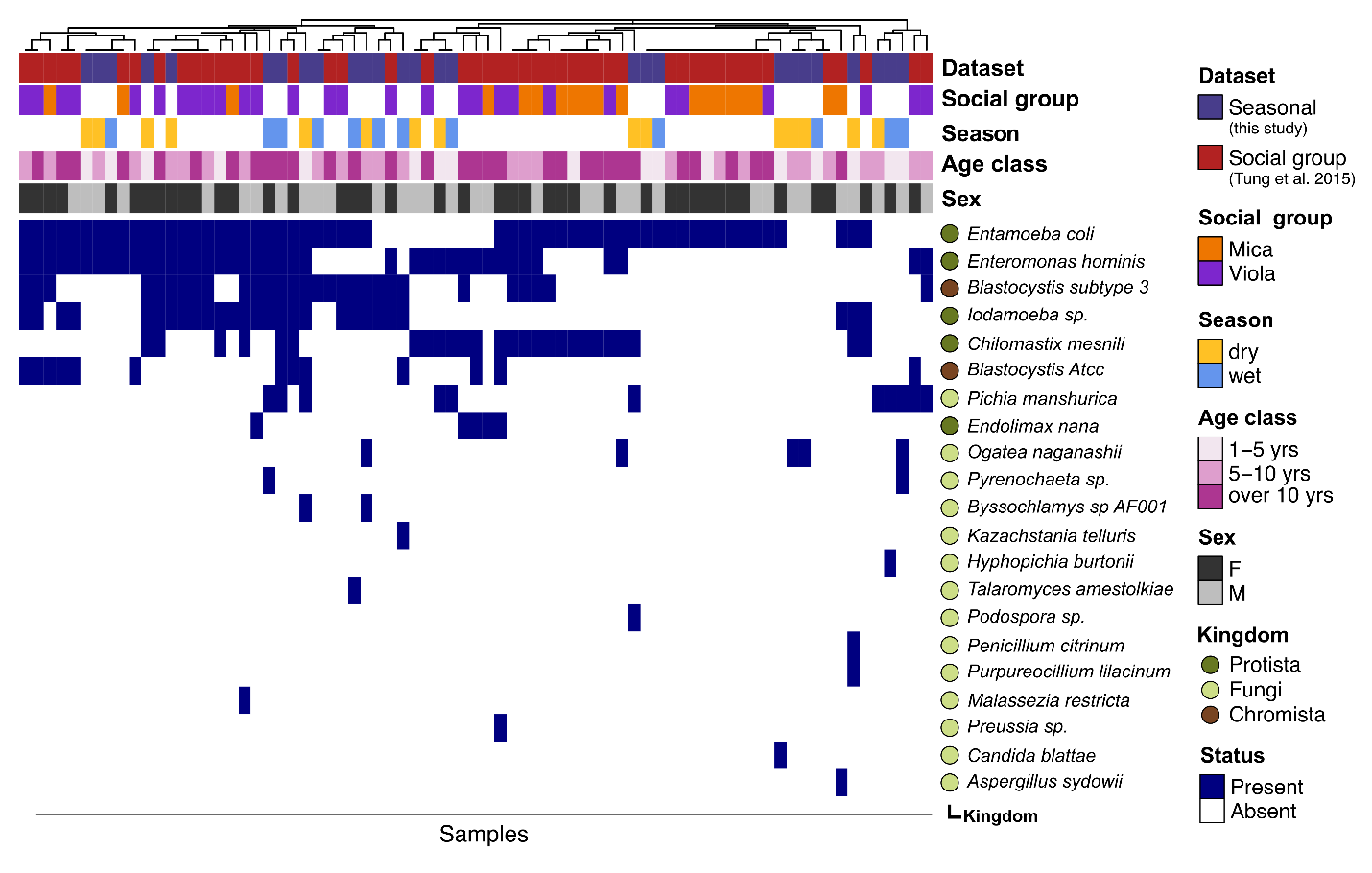

**Supplementary Figure 1. Eukaryotic diversity across both data sets.** Heatmap of the eukaryotic taxonomic composition across the two datasets. Each column represents one fecal sample, and the colored rows indicate, from top to bottom, data set, social group (n=48 samples), season (n=27 samples), age class, and sex of the host. Circles and names to the right of the heat map show the eukaryotic taxonomic classification. Colors for taxa circles indicate classification at the kingdom level. The sample clustering at the top of the heatmap reflects Euclidean distance of eukaryotic community composition.
